# Effects of the GABA_B_ receptor agonist 3-aminopropyl (methyl) phosphinic acid on food intake in free feeding rats

**DOI:** 10.64898/2026.01.08.698375

**Authors:** Ivor S Ebenezer

## Abstract

The GABA_B_ agonist baclofen has been shown to increase food intake in rats by a GABA_B_ receptor mediated mechanism. The present study, was undertaken to extend these observations by investigating the effects of intraperitoneal (ip) administration of the potent GABA_B_ receptor agonist 3-aminopropyl (methyl)phosphinic acid (SKF97541) on food intake in free feeding rats. In the first experiment, male Wistar rats (n=6) received intraperitoneal (ip) injections of saline or SKF97541 (0.1, 0.2, or 0.4 mg / kg), with food intake monitored over 120 minutes. The results showed that the 0.1 and 0.2 mg/kg doses of SKF97541 significantly increased cumulative food intake at 120 min (at least P<0.05) whereas the 0.4 mg/kg dose had no acute effects on feeding. However, the 0.4 mg / kg dose produced ataxia and sedation in the rats and these effects may have competed with feeding behaviours. A second experiment was carried out to investigate this possibility. Rats (n=6 in each of two groups) received daily injections of either saline or SKF97541 (0.4 mg /kg, ip) for five days. By day 5, the rats developed tolerance to the sedative and ataxic effects, revealing a significant increase in food intake at 120 min compared to controls (P<0.05). The results obtained in this study with the potent GABA_B_ agonist SKF97541 confirm and extend previous observations with baclofen and provide further support for a role of GABA_B_ receptors in the regulation of food intake.

## 1. Introduction

Gamma-amino butyric acid (GABA) is the major inhibitory neurotransmitter in mammalian central nervous system. It acts at two distinct receptor subtypes, namely the ionotropic GABA_A_ and the metabotropic GABA_B_ receptor (Olsen, 2002; Bowery and Smart, 2006, Ebenezer, 2015). There is also a subtype of the GABA_A_ receptor with distinctive pharmacological properties, previously known as the GABA_C_ receptor, but is now referred to as the GABA_A-ρ_ receptor (Naffaa et al., 2017). GABA is found in high concentrations in areas considered to be involved in the control of feeding behaviour, such as the lateral, ventromedial and arcuate nuclei of the hypothalamus (Decavel and Van den Pol, 1990; Backberg et al., 2003) and it has been demonstrated that intracerebroventricular (icv) injections of the GABA_B_ receptor agonist baclofen elicited feeding in satiated pigs and non-deprived rats by an action at central GABA_B_ receptors (Ebenezer, 1990; Ebenezer and Baldwin, 1990) and that systemic administration of baclofen increases feeding in free feeding or satiated rats (Ebenezer and Pringle, 1992, Ebenezer, 1995; Ebenezer and Patel, 2004; Higgs and Barber, 2004; Buda-Levin et al., 2005; Patel and Ebenezer, 2008a) and mice (Ebenezer and Prabhaker, 2007). Experiments conducted in our laboratory have suggested that endogenous GABA, acting at central GABA_B_ receptors, plays a physiological role in the regulation of feeding behaviour (Patel and Ebenezer, 2004) and as baclofen readily crosses the blood brain barrier (Faigle and Keberle, 1972) that the hyperphagia produced by systemic administration of the drug is mediated by a central mechanism of action (see Ebenezer and Patel, 2004).

Pharmacological and immunohistochemical studies have implicated a number of brain areas, such as the medial raphe nucleus, the nucleus accumbens and the arcuate nucleus of the hypothalamus, as possible central sites where the drug may act to mediate its hyperphagic actions (Wirtshafter et al., 1993, Stratford and Kelley, 1997, Ward et al., 2000, Backberg et al., 2003).

While most of the studies have used baclofen to investigate the relationship between GABA_B_ receptors and food intake, there has been a marked paucity of studies using other GABA_B_ receptor agonists. In the present study, the dose-related effects of the potent GABA_B_ receptor agonist 3-aminopropyl (methyl) phosphinic acid (known by the research code SKF97541; Froestl et al., 1995; Lehman et al., 2011) was investigated in free feeding rats to assess if it elicited similar effects as those of baclofen on food intake.

## 2. Material and methods

The protocols used in this study were approved by the Ethical Review Committee at the University of Portsmouth, U.K. and carried out under licence granted by the United Kingdom Home Office.

### 2.1 The dose-related effects of 3-aminopropyl (methyl) phosphinic acid (SKF-97541) food intake in free feeding rats

Adult male Wister rats (n=6; body weights: 380 - 420g) were housed in cages in groups of 3 where they had free access to food and water at all times. The animals were handled regularly and maintained on a 12 h light/ dark cycle (lights on at 830 h and lights off at 2030 h). The rats were given four training sessions where they were allowed free access to their normal pelleted food (Food composition: (a) Percentage mass: protein 20%, oil 4.5%, carbohydrate 60%, fibre 5%, ash, 7% + traces of vitamins and metals, (b) Percentage energy: protein 27.3%, oil 11.48% and carbohydrate 61.2%, (c) Energy density: 3.600 kcal/g) and water in experimental cages measuring 32×25×10 cm.The food was presented to the rats in shallow cylindrical cups. During the experimental sessions that followed, each animal was injected intraperitoneally (ip) with either physiological saline (control) or SKF-97541 (0.1, 0.2 or 4.0 mg/kg) immediately before being placed in the experimental cage for 120 min with free access to food and water. The amount of food consumed for each rat was measured at 30, 60 and 120 min, as described previously (Ebenezer, 1990). A repeated measures design was used, with each rat receiving all doses of saline or SKF in a Latin square design. The animals had free access to water in the experimental cages. A period of 3 to 4 drug-free days was allowed between successive trials.

### 2.2. Effect of repeated administration of 3-aminopropyl (methyl) phosphinic acid (SKF-97541; 0.4 mg/kg) on food intake in free feeding rats

Adult male Wistar rats (n=12; body weights: 365 - 415) were divided into two equal groups and housed in cages in groups of 4. They received similar training sessions in experimental cages as described for Experiment 1. The rats were injected ip once daily for 5 days with either physiological saline solution (n=6) or with SKF-97541 (0.4 mg / kg; n=6). On Day 1 and Day 5 the rats were placed in the experiment cages for 2h with free access to food and water immediately after injection and food intake recorded as described for Experiment 1.

### 2.3 Drugs

3-Aminopropyl (methyl) phosphinic acid (SKF97541) was purchased from Sigma Biochemicals, Dorset, UK. The drug was dissolved in physiological saline solution (0.9% w/v, NaCl) to give an injection volume of 0.1 ml/100 g body weight. Physiological saline solution was used as the control.

### 2.4 Statistics

The cumulative food intake data at each measurement time point for Experiment 1 were analysed by one way analysis of variance (ANOVA) with repeated measures on treatment followed by the Tukey *post-hoc* test (Winer, 1971). The data for Experiment 2 at each measurement time point was analysed by two way ANOVA with repeated measures on treatment and time (Days) followed by the Tukey *post-hoc* test.

## 3. Results

### 3.1 The dose-related effects of 3-aminopropyl (methyl) phosphinic acid (SKF-97541 ) on food intake in free feeding rats

The effects of physiological saline and SKF (0.1, 0.2 and 0.4 mg /kg) on cumulative food intake at 30, 60 and 120 min are illustrated in Figure 1. Analysis of the data showed that there was no significant differences compared with control data on cumulative food intake at 30 and 60 min (F_(3,15)_ = 1.97, P= 0.17, NS at 30 min; F_(3,15)_ = 2.46, P=0.10, NS at 60 min). However, ANOVA revealed a significant effect of drug treatment compared with control data (F _(3,15)_ = 12.3, P<0.01) on cumulative food consumption at 120 min. *Post hoc* tests showed that the 0.1 and 0.2 mg / kg doses of SKF97541 significantly increased cumulative food intake at 120 min (P<0.05 and P<0.01 respectively).

**Figure 1.**
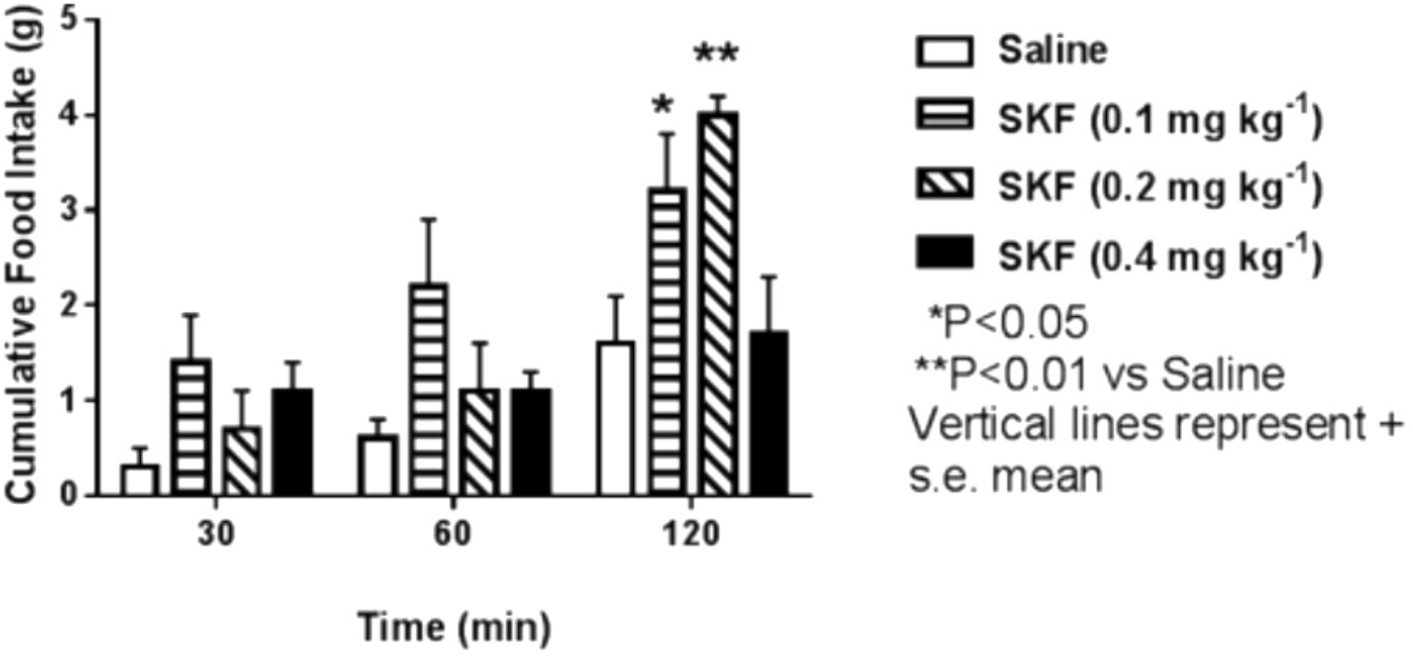
The dose related effects of SKF97541 (0.1, 0.2 and 0.4 mg / kg) on food intake in free feeding rats. Refer to the text for further details.

All doses of SKF produced signs of ataxia and mild sedation in the rats that were doses related in intensity. It was mainly apparent during the first 30 min for all doses and these effects could have competed with feeding behaviours.

### 3.2 Effect of repeated administration of 3-aminopropyl (methyl) phosphinic acid (SKF-97541) (0.4 mg/kg) on food intake in free feeding rats

The results of the second experiment experiment is shown in Table 1. Analysis of the cumulative food intake data at 30 and 60 min showed that there were no significant main effects of treatment (F_(1,10)_ = 3.08, P= 0.112 at 30 min; F_(1,10)_ = 1.90, P<0.198 at 60 min) but there were significant effects of time (Day) (F_(1,10)_ = 11.23, P<0.01 at 30 mins; F_(1,10)_ = 12.5, P<0.01 at 60 min). There were no significant effects of treatment x time interactions. By contrast, at 120 min, ANOVA revealed significant main effects of treatment (F_(1,10)_ = 4.98, P<0.05) and time (Days) (F_(1,10)_ = 22.15, P<0.01), but was just below the level of significance for treatment x time interaction (F_(1,10)_ = 4,67, P<0.055, NS). *Post hoc* tests showed that the food intake was (i) not significantly different between the saline control and SKF-97541 treated rats on Day 1 (P=0.3760, NS) and (ii) between the saline controls on Day 1 and Day 5 ( P= 0.136, NS). However, SKF-97541 significantly increased cumulative food intake compared with the saline control data at 120 min on Day 5 (P<0.05). Furthermore, the effect of SKF-97541on cumulative food intake at 120 min on Day 5 was significantly greater than that recorded at 120 min on Day 1 (P<0.01). Thus, while on Day 1, SKF97541 (0.4 mg / kg had no effect on food consumption, it significantly increased cumulative food intake on Day 5 after the animal had received 4 doses of the drug on the previous days.

**Table 1.** Effects of daily administration of SKF97541 (0.4 mg / kg) on food intake in rats measured on Day 1 and Day 5.

| Time (min) | Food Consumption (g) +/- SEM |  |  |  |
| --- | --- | --- | --- | --- |
|  | Day 1 |  | Day 5 |  |
|  | Saline | SKF | Saline | SKF |
| 0-30 | 0.66 +/- 0.2 | 0.7 +/- 0.3 | 1.1 +/- 0.3 | 1.9 +/- 0.3 |
| 0-60 | 1.1 +/- 0.2. | 1.2 +/- 0.2 | 1.6 +/- 0.5 | 2.4 +/- 0.4 |
| 0-120 | 1.4 +/- 0.3 | 1.7 +/- 0.4 | 2.4 +/- 0.4 | 4.4 +/- 0.8*# |

\*P<0.05 compared with Saline at 120 min on Day 5 #P<0.01 compared with SKF97541 at 120 min on Day 1

SKF97541 (0.4 mg /kg) produced ataxia and mild sedation in the rats on Day 1 that was most apparent during the first 30 min during the trial in the experimental cages. However, on Day 5, there was a marked attenuation of these effects after injection of SKF97541 compared with those observed on Day 1.

## 4. Discussion

3-Aminopropyl(methyl)phosphinic acid (SKF97541) is a potent agonist at GABA_B_ receptors (Seabrook et al., 1990, Froestl et al, 1995, Lehmann et al, 2011). Current estimates suggest it is at least ten times more potent than baclofen (Piqueras and Martinez, 2004). Consequently, the present study utilised doses of SKF97541 ten times lower than the baclofen dosages previously shown to increase food intake (Ebenezer and Pringle, 1992, Ebenezer, 1995, Patel and Ebenezer, 2008 a,b, Ebenezer and Patel, 2011). Although SKF97541 has been reported to possess weak antagonist activity at the GABA_A-ρ_ receptor (formerly designated as the GABAC receptor; Naffaa et al., 2017), these receptors are primarily localised in the retina (Naffaa et al., 2017) and it is therefore unlikely that this action of SKF97541 will influence feeding behaviour.

The results of Experiment 1 (see Fig. 1) confirm previous findings obtained with baclofen (Ebenezer and Pringle, 1992, Ebenezer, 1995; Ebenezer and Patel, 2004; Higgs and Barber, 2004; Buda-Levin et al., 2005; Patel and Ebenezer, 2008a, Ebenezer and Patel, 2011) and show that the lower doses of SKF97541 (0.1 and 0.2 mg / kg) significantly increase food intake in free-feeding rats. This hyperphagic effect was most pronounced 120 minutes post-administration; no significant increases were observed at the 30 min or 60 min intervals. It is notable that while ANOVA indicated no significant treatment effects at these earlier time points, the mean food intake for the 0.1 mg/kg dose remained higher than control values (Fig. 1).

By contrast, the 0.4 mg/kg dose did not significantly affect food intake. However, all doses of SKF97541 induced signs of ataxia and sedation, particularly within the first 30 minutes. These depressant effects were dose-dependent, reaching peak intensity at the 0.4 mg/kg dose. Similar sedative properties have been reported in mice, where a 0.3 mg / kg dose (ip) decreased locomotor activity and potentiated ethanol-induced sedation (Besheer et al., 2004). It is therefore highly probable that these side effects competed with and masked the drug’s orexigenic effects at the higher dose.

Previous research has indicated that moderate to high systemic doses of baclofen can cause ataxia and sedation that interfere with feeding-related behaviours (Ebenezer and Pringle, 1992, Patel and Ebenezer, 2010, Ebenezer and Patel, 2011, Bains and Ebenezer, 2013.). However, while tolerance typically develops to these depressant effects upon repeated administration, tolerance does not develop to the hyperphagic effects of the drug (Patel and Ebenezer, 2008 a,b). Experiment 2 was therefore undertaken to investigate whether SKF97541 exhibited a similar profile with repeated daily injections of 0.4 mg / kg. The results demonstrate that after five days of treatment, the rats developed tolerance to the drug’s depressant effects. Consequently, a significant increase in food intake was observed 120 minutes post-administration, revealing the underlying hyperphagic effect of the 0.4 mg/kg dose once the sedative interference was diminished.

The primary finding of this study is that SKF97541 acts as a highly potent stimulator of food intake, achieving hyperphagic effects at roughly 10% of the dose required for baclofen. This aligns with the known high affinity of SKF97541 for the metabotropic GABA_B_ receptor. A critical challenge in studying GABAergic influences on food intake is the overlap between orexigenic signals and motor suppression. The results with the 0.4 mg/kg dose highlight this phenomenon of competing behaviours. The drug’s sedative and ataxic effects initially dominate, preventing the animal from engaging in the motor tasks required for feeding. As the sedative effects wane (either through metabolism with the lower doses in Experiment 1 or development of tolerance in Experiment 2), the underlying orexigenic signal elicited by the drug becomes apparent.

In summary, these findings confirm and extend previous observations of GABA_B_ receptor mediated feeding behaviour. The results demonstrate that the GABA_B_ agonist SKF97541 effectively increases food intake in free-feeding rats, mirroring the effects of baclofen but at significantly lower doses because of its higher potency.

